# Hepatitis E virus infection upregulates ING5 expression in *vitro* and *vivo*

**DOI:** 10.1101/2024.03.22.586368

**Authors:** Wanqiu Zhao, Yueping Xia, Tengyuan Li, Huichan Liu, Guo Zhong, Dongxue Chen, Wenhai Yu, Yunlong Li, Fen Huang

## Abstract

Hepatitis E virus (HEV) is the major pathogen of viral hepatitis. Immunocompromised individual infected by HEV is prone to chronic hepatitis and increases the risk of hepato-cellular carcinoma (HCC). Inhibitor of growth family member 5 (ING5) is a tumor suppressor gene, is low expressed in cancer tumor or cells. However, the underlying relationship between ING5 and HEV infection is unclear. In the present study acute and chronic HEV animal models were employed to explore the interaction between ING5 and HEV. Notably, the expression of ING5 was significantly increased both in the liver of acute HEV infected BALB/c mice and chronic HEV infected rhesus macaques. In addition, the relationship between HEV infection and ING5 expression was further identified in human hepatoma (HepG-2) cells. In conclusion, HEV infection strongly upregulated ING5 expression in *vivo* and in *vitro*, bearing significant implications in further understanding the pathogenic mechanism of HEV infection.

## 1. Introduction

Hepatitis E virus (HEV) is the most common cause of acute viral hepatitis worldwide [1]. HEV infection usually causes acute self-limiting diseases with low mortality in general population, but leads to about 25% maternal death in pregnant women [2-4]. Importantly, chronic HEV infection had been reported in immunocompromised patients, such as organ-transplant recipients [5, 6], HIV infected patients and cancer patients receiving chemotherapy [7]. HEV infection increases the risk of liver decompensation in patients with underling chronic liver disease, including non-alcoholic fatty liver disease (NAFLD), alcoholic liver disease (ALD) and chronic hepatitis C virus (HCV) [8-12] and aggravates the development of hepato-cellular carcinoma (HCC) in chronic HBV infected patients. However, the pathogenesis of HEV infection promote the development of HCC is still unclear.

Inhibitor of growth (ING) family (member 1-5) play important roles in cell cycle, cell proliferation and growth, apoptosis and senescence [13, 14]. The ING family has been identified to be a tumor suppressor gene (TSG), interacted with the determinants of chromatin function and gene-specific transcription factors [13]. ING5 encodes a tumor suppressor protein that inhibits cell growth and induces apoptosis [13]. It contains a PHD-type zinc finger and interacts with tumor suppressor p53 and p300, a component of the histone acetyltransferase complex, indicating a role in transcriptional regulation [15, 16]. ING5 plays a crucial role in chromatin acetylation, oncogenic transformation and cancer development [16]. In cancer cells, ING5 transcript levels are often suppressed, for example lowly expressed ING5 was found in esophageal squamous cell carcinoma (ESCC) [17], and suppressed ING5 reported in those HBV induced HCC [18, 19]. The mechanism of suppression of ING5 expression may have to do with the abnormally high methylation levels of the ING gene promoter, which have been correlated with low transcript levels [13]. Thus, ING5 is considered to be a promising target for cancer treatment.

HEV infection in immunocompromised patients can lead to chronic hepatitis and progress to cirrhosis [20, 21]. However, whether there is a regulation of ING5 during HEV infection is still unknown. In the present study, the expression of ING5 in acute HEV infected animal model and chronic HEV infected animal model were assessed to explore the potential regulation between HEV and ING5, providing novel insights into the pathogenic mechanisms of HEV infection.

## 2. Materials and methods

### 2.1 HEV Strain and Cells

Hepatoma cells (HepG-2) were maintained in DMEM supplemented with 10% FBS at 37 °C under 5% CO_2_. HepG-2 cells were infected with HEV (KM01 strain, KJ155502, isolated from a HEV-positive stool sample in Kunming City, genotype 4, 2 × 10^6^ copies/mL) in accordance with our previous studies [22, 23]. The cells were collected at 48 and 72 h post-infection (hpi) for viral quantification by quantitative real-time PCR (qRT-PCR) or fixed at 72 hpi for immunofluorescence assay (IFA).

### 2.2 Reagents and animal tissues

HEV-specific antibody (ab233244), HRP-labeled secondary antibody (ab205718), and DAB Substrate Kit (ab64238) were purchased from Abcam (Cambridge, UK). Rabbit-anti-ING5 (10665-1-AP) was purchased from Proteintech (Boston, MA, USA). HRP-, TRITC-, and FITC-labeled secondary antibodies (AS026, AS011) were purchased from ABclonal (Boston, MA, USA). Immobilon ECL Kit (WBKLS0500) was purchased from Milillpore (Billerica, USA). Trizol reagent (15596-026) was purchased from (Invitrogen, USA). M-MLV reverse transcriptase (2641A) and TB Green^®^ Fast qPCR Mix (RR430A) were purchased from Takara (Tokyo, Japan).

The animal tissues used in this study were collected from our previous studies [24, 25]. In a brief, the liver of rhesus macaques with or without HEV infection at 272, 650, and 770 days post infected (dpi) and the liver and ovary of BALB/c mice with or without HEV infection at 7, 14, 21, 28, and 36 dpi were collected. All animal experiments were preserved by the Laboratory of Viral Infection and Immunology of School of Medicine in Kunming University of Science and Technology.

### 2.3 RNA isolation and qRT-PCR

Total RNA was extracted from tissue by using Trizol reagent in accordance with the manufacturer’s instructions. Reverse transcription was performed using M-MLV reverse transcriptase. RNA expression levels were quantified by TB Green-based qRT-PCR with the primers of ING5 sense 5′-CTTCATCCAAAACGATGATGC-3′ and ING5 antisense 5′-CGTTCCAAAAAATACTTTATT-3′. HEV RNA was quantified as described in previous study [26]. qRT-PCR was performed in a Bio-Rad CFX96TM Real-Time PCR System. GAPDH served as a loading control. The relative gene expression was calculated by the 2^−ΔΔCT^ method. qRT-PCR was performed in the Bio-Rad CFX96TM Real-Time PCR System.

### 2.4 Western blotting assay

Cells and tissues were harvested and lysed with RIPA buffer. An equivalent amount of total protein was separated through 10% sodium dodecyl sulfate–polyacrylamide gel electrophoresis and then transferred onto a nitrocellulose membrane. Nonspecific binding sites were blocked with 5% skim milk, and the membrane was incubated overnight separately with primary antibodies (ING5 1:1000 dilution and GAPDH 1:5000 dilution). HRP‐conjugated IgG was used as a secondary antibody (1:5000 dilution). GAPDH served as the loading control. The bands were exposed to X‐ray films by using an Immobilon ECL Kit.

### 2.5 Immunohistochemistry (IHC) assay

Tissue samples were fixed in 10% neutral-buffered formalin and embedded in paraffin. Specimens were cut into 4 µm serial sections. Standard hematoxylin staining was performed, and tissues were examined under a microscope. Tissues were deparaffinized, hydrated, heated in a water bath for antigen retrieval, and blocked with the addition of 3% hydrogen peroxide for 15 min. Tissue sections were incubated with the primary antibodies (ING5 1:100 dilution and HEV 1:200 dilution) overnight at 4 °C. The sections were washed with PBS three times and incubated with HRP-labeled secondary antibodies at 37 °C for 1 h. Nuclei were counterstained with DAB substrate and viewed under a microscope (Nikon, E200, Japan).

### 2.6 Immunofluorescence assay (IFA)

Cells were washed three times with PBS after 48 or 72 hpi, fixed with 4% paraformaldehyde for 10 min, and washed with PBS. Cells were treated with primary antibodies (ING5 1:100 dilution and 5F3 1:200 dilution) and incubated with HRP-labeled secondary antibodies at 37 °C for 1 h. Nuclei were stained with DAPI and observed under a confocal fluorescence microscope (Nikon, Ts2, Japan).

### 2.7 Statistical analysis

This study used GraphPad Prism 9 and Image-Pro-Plus 6.0 for statistical analysis. The mean ± standard deviation was used for the normal -distribution expression measurement data, and two groups were compared using *Student’s* t-test. We represented the non-normal distribution measurement data by the median (quartile), and two groups were compared using the Mann-Whitney U test. A P value <0.05 was indicated to indicate statistically significant differences.

## 3. Results

### 3.1 Acute HEV infection significantly elevated ING5 expression in *vivo*

HEV infects BALB/c mice and induces acute infection symptoms had been confirmed in our previous study [25], for instance, acute hepatitis E and viral genome are detectable for about 4 weeks. Accordingly, using this HEV infection model, we found that ING5 protein expression significantly increased upon HEV infection for 7, 14, 21, 28, or 36 days after evaluating the liver tissues by IHC (Figure 1A). The HEV and ING5 protein expression levels were quantified by grayscale analysis (Figure 1B). Interestingly, the expression level of ING5 and HEV was found to have a similar tendency, indicating that a reciprocal interaction between ING5 expression and HEV infection. For example, the ING5 protein level was stably upregulated throughout the 3 weeks but was only slightly downregulated at 28 dpi. A similar trend was observed in HEV replication. However, no positive signal of the HEV ORF2 were detected at 36 dpi indicated that the viruses were cleared, and the ING5 expression consistently decreased. Subsequently, the expression of HEV RNA and ING5 at the mRNA and protein levels were quantificated by qRT-PCR or Western blotting in the liver. Notably, the genes of HEV RNA and ING5 and also proteins levels in the liver still increased at 36 dpi even after the virus was cleared (Figure 1E and 1F).

**Figure 1.**
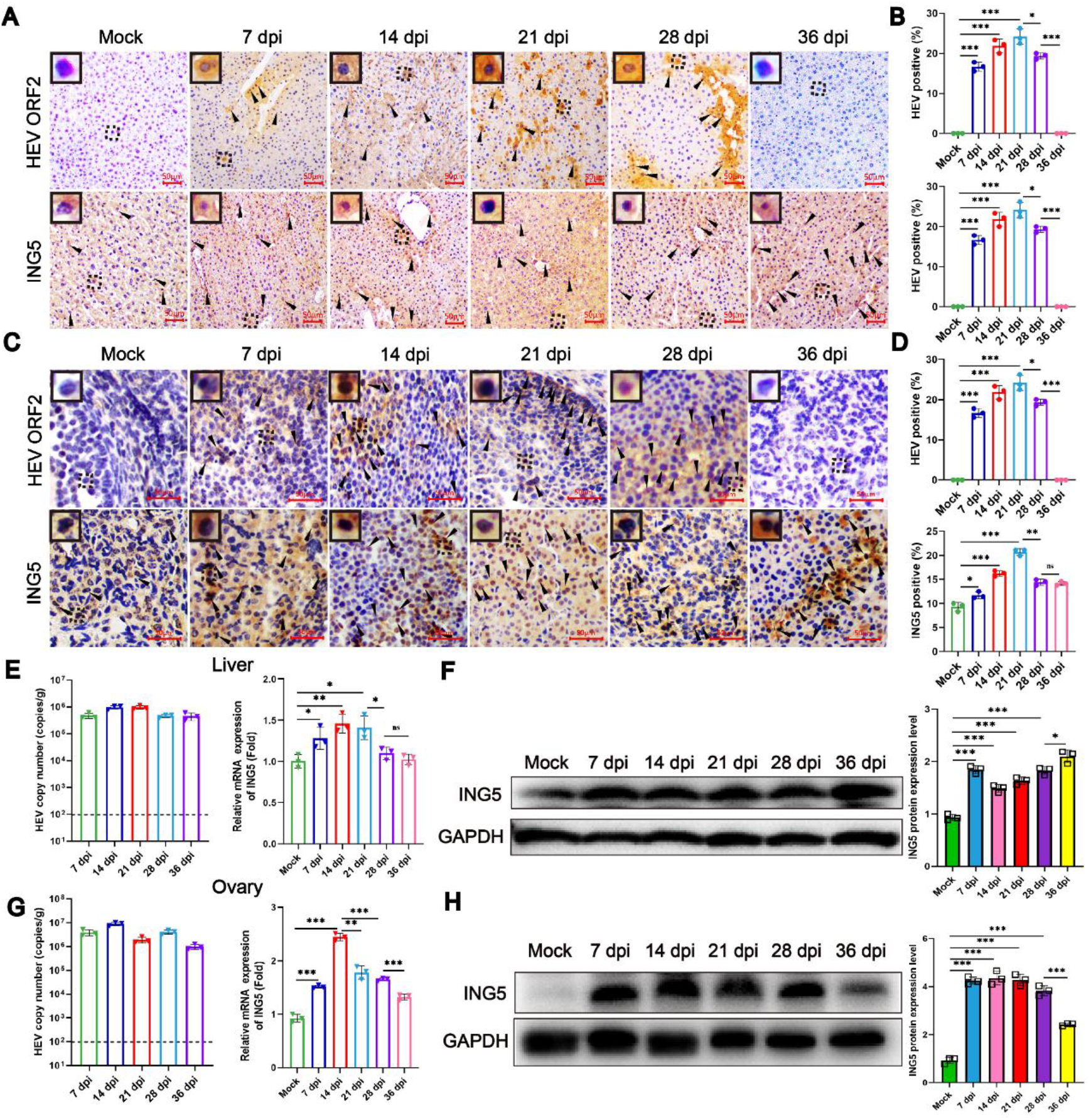
Acute HEV infection activates ING5 high expression in BALB/c mice liver and ovary tissues. **(A)** BALB/c mice were infected with HEV for different times. IHC showed HEV ORF2 in infected, but not in the control, animal liver tissue, and ING5 higher expression in the infected group. Scale bar, 50 μm. **(B)** The positive signaling of HEV particle and ING5 protein in liver tissue was measured at 7, 14, 21, 28, or 36 dpi (n = 3). **(C)** IHC-analyzed HEV infection and ING5 expression in the ovary tissues. Scale bar, 50 μm. **(D)** The specific levels of HEV infection and ING5 expression in ovaries were quantified and analyzed (n = 3). **(E)** HEV RNA and ING5 mRNA levels were quantified by qRT-PCR of liver tissues (n = 3). **(F)** ING5 protein in liver lysates was determined by Western blotting and quantified by gradation analysis (n = 3). **(G)** HEV RNA and ING5 mRNA levels were quantified by qRT-PCR in the ovary tissues (n = 3). **(H)** ING5 protein in ovary lysates was determined by Western blotting and quantified by gradation analysis (n = 3). Data are expressed as the mean ± SD. *p < 0.05; **p < 0.01; ***p < 0.001; ****p < 0.0001.

HEV has multiple extra-hepatic replication sites, including placenta, uterus, and ovary [25, 27, 28]. Accordingly, we further profiled the dynamics of ING5 protein expression in ovary tissues upon HEV infection. Increased ING5 and HEV were observed over time (Figure 1C and 1D). Expectedly, a similar tendency of ING5 activation and HEV replication was observed in liver and ovary tissues. The expression of HEV RNA and ING5 in the ovary tissues to further confirm the potential relationship between HEV infection and ING5 activation (Figure 1H and 1G). All results showed a dynamic coordination of ING5 expression and HEV infection in the BALB/c mice model.

### 3.2 Chronic HEV infection potently increased ING5 expression in *vivo*

Chronic HEV infection has been identified in various patients, including recipients of solid-organ transplantation, those receiving immunosuppressive therapies, HIV co-infected, and with liver disease. Persistent HEV infection has a high probability of causing liver fibrosis and rapidly progressing to cirrhosis. Thus, the potential interaction between ING5 regulation and HEV infection in chronic HEV-infected rhesus macaques were further investigated. Notably, the ING5 protein had stably upregulated expression throughout the chronic HEV infection period (up to 770 dpi) (Figure 2A). Subsequently, the level of ING5 protein was calculated (Figure 2B). The increased HEV RNA and ING5 mRNA and protein expression levels, further confirmed the potential cooperation between HEV infection and ING5 activation in HEV chronic infected animal model (Figure 2C and 2D), which is consistent with that in HEV-infected BALB/c mice. Notably, HEV infection activated the expression of ING5 in both acute and chronic infection.

**Figure 2.**
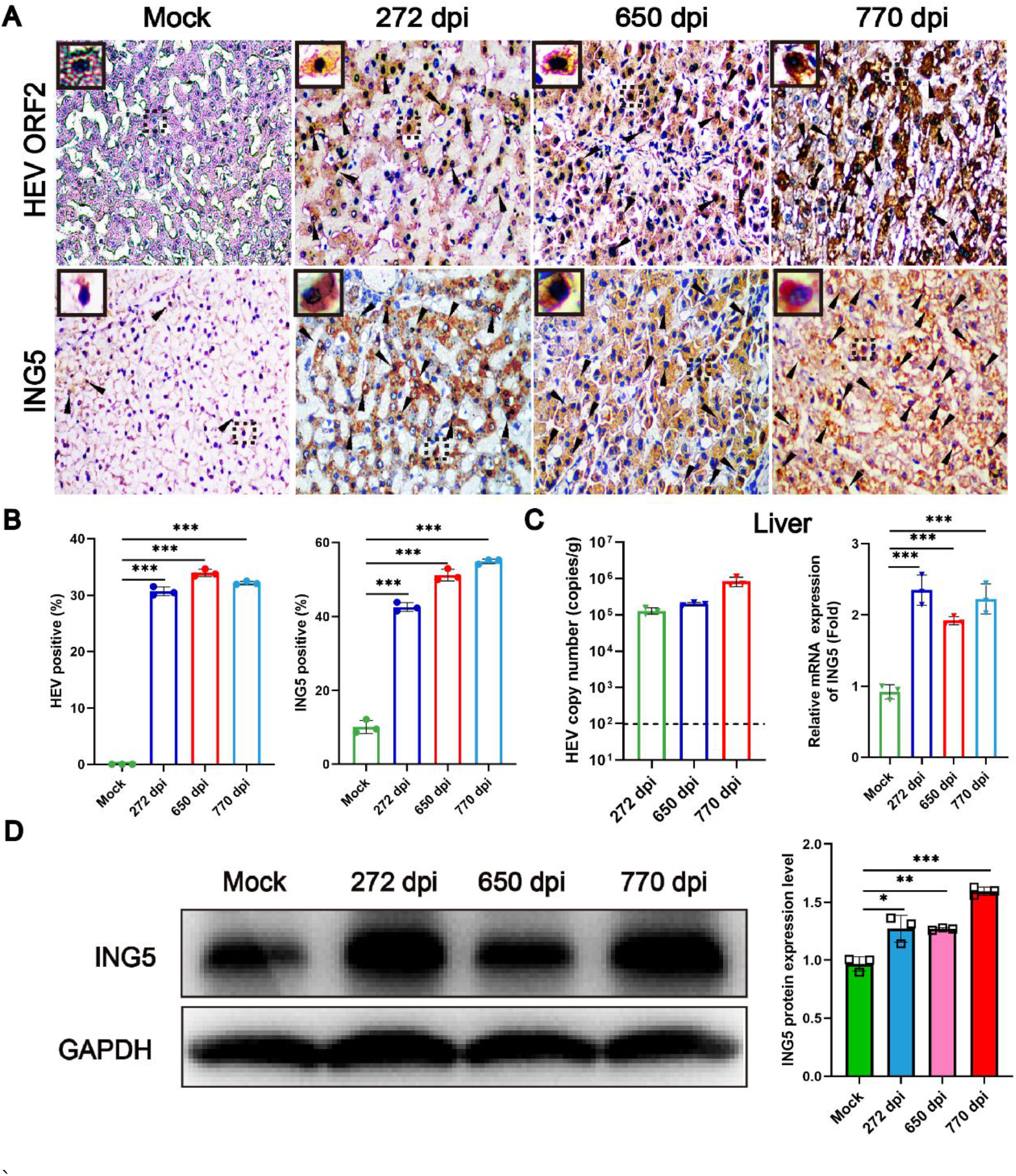
ING5 robust expression in the liver tissue of chronic HEV-infected rhesus macaques. **(A)** IHC showing HEV ORF2 and ING5 protein expression in the liver tissues of chronic HEV-infected rhesus macaques at 272, 650, or 770 dpi. Scale bar, 20 μm. **(B)** Quantification of the positive signaling of HEV ORF2 and ING5 protein in liver tissues (n = 3). **(C)** HEV RNA and ING5 mRNA levels of liver tissues quantified by qRT-PCR (n = 3). **(D)** ING5 protein in liver lysates determined by Western blotting and quantified by grayscale analysis (n = 3). Data are the mean ± SD. *p < 0.05; **p < 0.01; ***p < 0.001; ****p < 0.0001.

### 3.3 Acute HEV infection robustly activated ING5 expression in *vitro*

Given the interaction between ING5 upregulated expression and HEV infection in BALB/c mice and rhesus macaques, we subsequently investigated the expression of ING5 in HEV-infected HepG-2 cells. First, HEV RNA and protein were detected by qRT-PCR and IFA to confirm viral replication after inoculating with infectious virus. HEV RNA and antigens were detected at 12-72 hpi (Figure 3A and 3C). Second, ING5 mRNA and protein expression levels were quantified by qRT-PCR, Western blotting, and IFA at the same time points and observed a similar tendency (Figure 3A, 3B, and 3C). In conclusion, HEV replication positive regulated ING5 expression.

**Figure 3.**
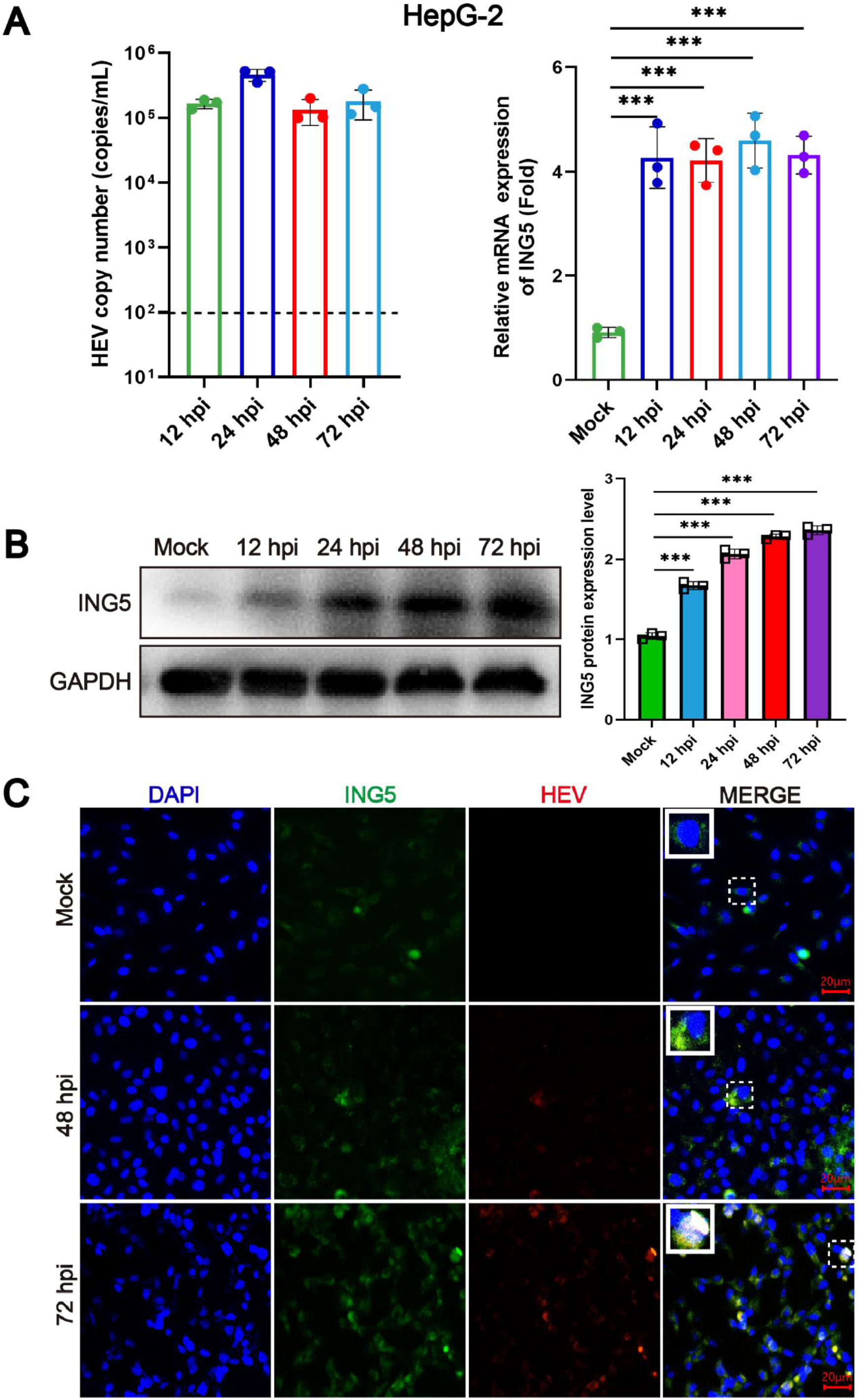
HEV-infected HepG-2 cells increased ING5 expression. **(A)** HEV RNA and ING5 mRNA levels were quantified by qRT-PCR (n = 3) in HEV-infected HepG-2 cells. **(B)** HepG-2 cells were infected with HEV for 12, 24, 48, and 72 h. ING5 protein in lysates was determined by Western blotting and quantified by gradation analysis (n = 3). **(C)** HepG-2 cells were infected with HEV for 48 and 72 h. Co-localization of ING5 (green) and HEV ORF2 (red) were examined under a confocal microscope. Nucleus was stained with DAPI (blue). Scale bar, 20 μm; 40× oil immersion objective. Data are the mean ± SD. *p < 0.05; **p < 0.01; ***p < 0.001; ****p < 0.0001.

### 3.4 Inhibition of HEV infection suppressed ING5 activation

Based on the results that HEV infection activated ING5 expression based on in *vivo* and in *vitro* models, the degree of HEV infection was positively correlated with ING5 expression. To further confirm this interaction, inhibition of HEV infection by using a classical inhibitor interferon α (IFN-α) to analyze its effect on ING5 expression. The mRNA and protein expression of ING5 upon HEV inhibition through qRT-PCR and Western blotting analyses were performed. Results showed that HEV was significantly suppressed by IFN-α, and the expression of ING5 decreased accordingly (Figure 4A and 4B), consistent with the reduction in the acute and chronic animal models. Overall, these results compellingly demonstrated that the expression of ING5 was influenced by HEV infection.

**Figure 4.**
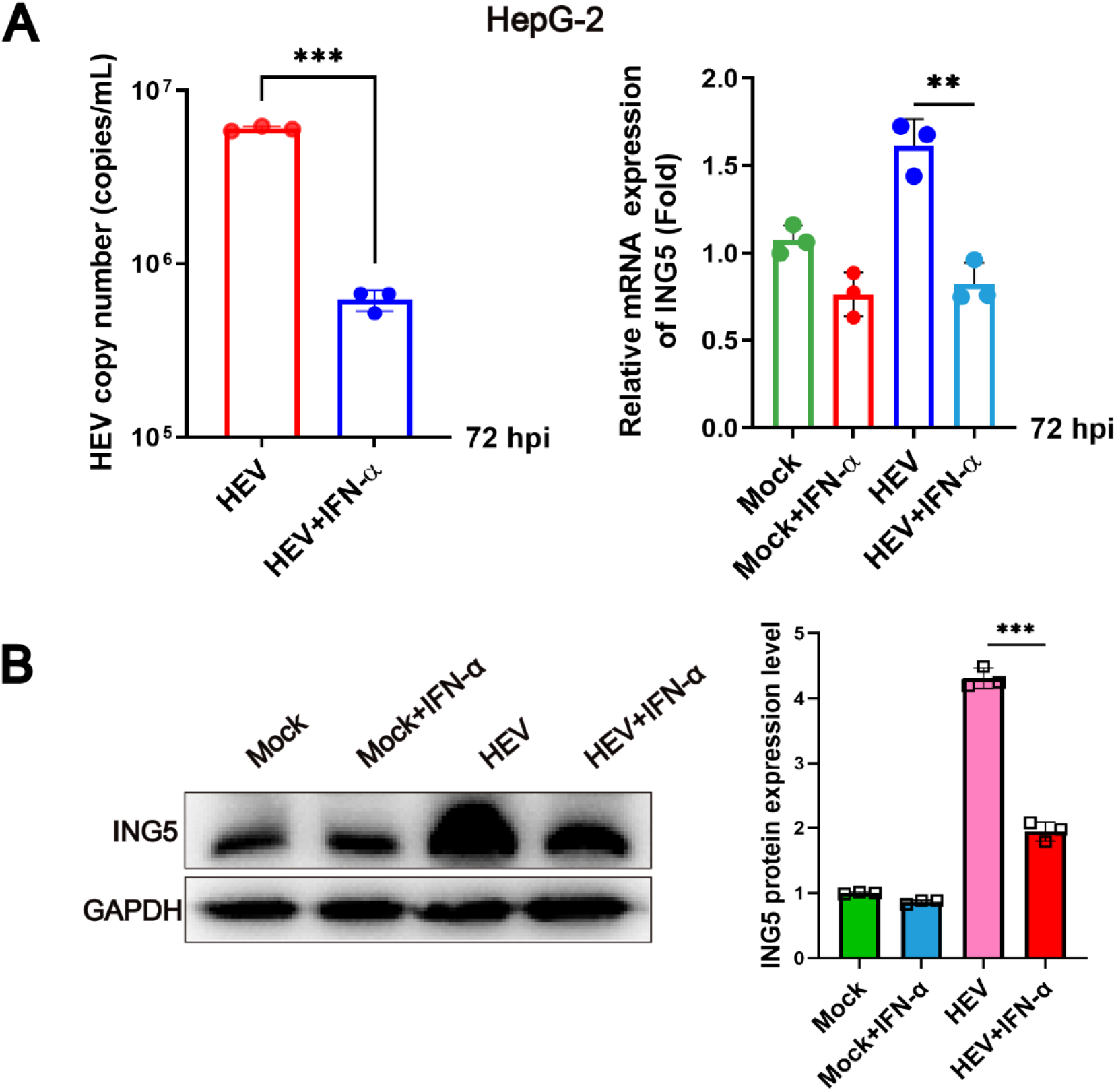
Inhibition of HEV replication abolished HEV-induced ING5 expression. HepG-2 cells were infected with HEV for 24 h and then treated with IFN-α (500 IU/mL) for 72 h. **(A)** HEV RNA and ING5 mRNA levels were quantified by qRT-PCR (n = 3). The uninfected group was set as the negative control. **(B)** ING5 protein levels were determined by Western blotting and quantified by grayscale analysis (n = 3). The uninfected group was set as the negative control (set as 1). Data are the mean ± SD. *p < 0.05; **p < 0.01; ***p < 0.001; ****p < 0.0001.

## 4. Discussion

HEV infection was thought to be an acute infection, but chronic HEV infection is widely reported in immunecomprised patient with a rapid progression to liver cirrhosis[21]. However, the pathogenesis of HEV induced liver cirrhosis is unexplored. In the present study, significantly activated ING5 was observed either in acute HEV infected BALB/c mice, or chronic HEV infected rhesus macaques. Although suppressed ING5 expression was found in HBV induced HCC, increased ING5 was determined in HEV infected human HCC cell lines (HepG-2), which indicated there is interaction between HEV infection and ING5 expression.

HEV infection causes a self-limiting disease, but acute liver failure or fulminant hepatic failure were occurred in HEV-infected pregnant women [3, 4]. HEV infection is associated with chronic hepatitis and liver cirrhosis, and increases the risk of HCC [29]. However, the relationship between HEV and HCC was rarely reported, and the role of HEV in developing HCC is unclear. HEV infection activates severe inflammatory responses in the liver [30], the long-term persistent inflammatory stimuli aggravates the liver damages, which may contribute to liver fibrosis, then cirrhosis and finally HCC, if no effective treatment. Furthermore, whether there is an association between the increased apoptosis in the liver of HEV infected animals/patients and the up-regulated ING5-related to HEV infection is need to be explored.

ING5 is a newly identified TSG. ING5 has been reported in lung cancer tumors to inhibit the ability of cancer cells to invade normal tumor-adjacent tissues and distant metastasis through the AKT pathway [31]. ING5 overexpression in the colorectal cancer cells suppresses the cells proliferation, migration, invasion, and promoted cell apoptosis, which could be reversed by ING5 knockdown [32]. Similarly, a negative correlation was found between the expression of ING5 and cell proliferation/migration in ovarian cancer cells [33]. ING5 inhibits migration and invasion of esophageal cancer cells by downregulating the IL-6/CXCL12 signaling pathway [31]. In addition, HBV infection suppresses ING5 to promote the proliferation of HCC cells [18]. ING5 is a potential molecular marker for carcinogenesis and following progression, and as a target for gene therapy. However, the expression of ING5 significantly activated in HEV infected acute BALB/c mice model and chronic rhesus macaques model, as well as in human hepatoma cells (HepG2). The increase of ING5 caused by HEV infection may play a critical role in the development of HCC, and more attentions should be payed. For example, the acetylated proteins in the liver should be further identified and explore their functions, which will benefit the knowledge of HEV pathogenesis.

## 5. Conclusion

HEV infection activates ING5 expression *in vivo* (BALB/c mice and rhesus macaques) and *in vitro* (HepG-2). Results demonstrated that HEV infection was positive regulates ING5 expression, and more attention should be payed to explore the potential regulatory mechanism, which will provide a new light for understanding the pathogenesis of HEV infection.

## Data availability

Data will be made available upon request.

## Ethical approval

The animal experimental protocol used in this study was approved by the Animal Protection and Use Committee of Kunming University of Science and Technology.

